# Robust Multiplicative Control in Chemical Reaction Networks Extended Version

**DOI:** 10.64898/2026.03.15.711890

**Authors:** Emmanouil Alexis, Clarence W. Rowley, José L. Avalos

**Affiliations:** Omenn-Darling Bioengineering Institute and the Department of Chemical and Biological Engineering, Princeton University, NJ, USA; Department of Mechanical and Aerospace Engineering, Princeton University, NJ, USA

**Author notes:** E- mail.

## Abstract

Achieving complex multi-species control objectives is essential for engineering advanced autoregulated biomolecular devices. This paper addresses the problem of robust steady-state tracking for outputs defined as multiplicative combinations of biomolecular species concentrations. We first introduce a control architecture realized via chemical reaction networks that steers the product of two target species concentrations in the controlled network to a prescribed value. A robust stability analysis is provided for closed-loop system families with distinct structural characteristics. The proposed framework is also extended to a more general formulation capable of regulating arbitrary monomial outputs involving multiple species. Numerical simulations of representative examples corroborate the theoretical results and illustrate the effectiveness of our approach.

## I. INTRODUCTION

Assembling biomolecular devices with user-defined properties lies at the heart of synthetic biology and has a profound impact across many real-world applications, ranging from molecular computing and robotics to living biotherapeutics and materials [1]–[5]. Achieving this requires the challenging task of designing chemical reaction networks (CRNs) with robust and predictable behavior intended to operate in noisy and uncertain biological environments [6]. This need has fueled the development of control-theoretic concepts tailored to biomolecular systems, including various feedback and feedforward control schemes realized via CRNs in *in vivo* and/or *in vitro* settings [7]–[10].

The vast majority of proposed CRN-based controllers have focused on regulating a single physical quantity, such as the concentration of a protein. Despite their success across a broad spectrum of practical applications, constructing autoregulated devices capable of reliably performing more sophisticated tasks necessitates the satisfaction of more complex control objectives involving multiple physical quantities of interest. Notably, a variety of centralized and decentralized regulation strategies for multi-output biomolecular processes are introduced in [11]. The main results of this study concern processes with two outputs of interest coupled through internal interactions, and demonstrate how their ratio, a linear combination of the outputs, or each output individually can be robustly controlled. Ratio-based regulation strategies tailored to different biological settings have also been investigated in [12]–[16]. In parallel, a two-input signal-processing module capable of ratiometric computation is implemented in [17].

Another interesting category of CRN-based controllers is multiplicative regulators. Although briefly discussed in [11], this direction remains largely unexplored. We refer to multiplicative regulation as the control of an output signal proportional to the (mathematical) product of multiple target physical quantities (e.g., species concentrations of interest), each possibly raised to a given power. In this work, we aim to fill this gap by introducing and analyzing a family of biomolecular control schemes suitable for this purpose. These schemes employ a centralized control approach based on integral feedback action, enabling robust steady-state tracking or, in biological terms, *robust perfect adaptation* (*RPA*). This property is realized through a family of biomolecular topologies that can be viewed as extensions of the *antithetic integral motif*, originally introduced in [18] and first experimentally implemented in [19].

We consider a biomolecular process to be controlled with partially known structure and subject to disturbances. We first introduce a structurally minimal class of CRNs capable of regulating the product of two target species concentrations, which represents the simplest multiplicative control objective. A key requirement for the proper functioning of the resulting closed-loop systems is asymptotic stability, which we analyze in depth for two structurally distinct system families, one of which encompasses the other as a special case. Our theoretical results are illustrated through two representative case studies. We further present a third case study involving a more complex control objective, together with a generalized control framework that enables regulation of outputs expressed as arbitrary monomials of multiple target species concentrations.

The remainder of this paper is organized as follows. Section II presents the notation and methodological approach adopted throughout this work. Section III introduces a control architecture of minimal structural complexity. Section IV provides a robust stability analysis of the resulting closed-loop systems. Section V discusses a practical application. Section VI discusses a generalized control formulation. Section VII summarizes the main contributions and outlines directions for future research.

## II. BACKGROUND

We consider chemical reactions [11], [20], [21] obeying the deterministic law of mass action, unless otherwise stated. In such cases (e.g., Hill-type kinetics), reaction rates are assumed to be continuously differentiable on their domain. Under mass-action kinetics, reactions can be broadly classified into three categories: (i) general transformation of reactants into products, (ii) catalytic production, and (iii) catalytic inhibition. Let *A* and *B* denote two biochemical species. These reaction types can be written in the generic forms A → *B*, A → A + *B*, and A + *B* → A, respectively. Their corresponding graphical representations are shown in Fig. 1 a. Furthermore, we model the dynamics of the chemical reaction networks (CRNs) under consideration using deterministic ordinary differential equations (ODEs) and perform all simulations in MATLAB (MathWorks).We assume that all species concentrations are finite and non-negative and all reaction rate constants are finite and positive (unless otherwise stated). For simplicity, each state variable denoting a species concentration is represented by the same uppercase letter as the corresponding species.

**Fig. 1:**
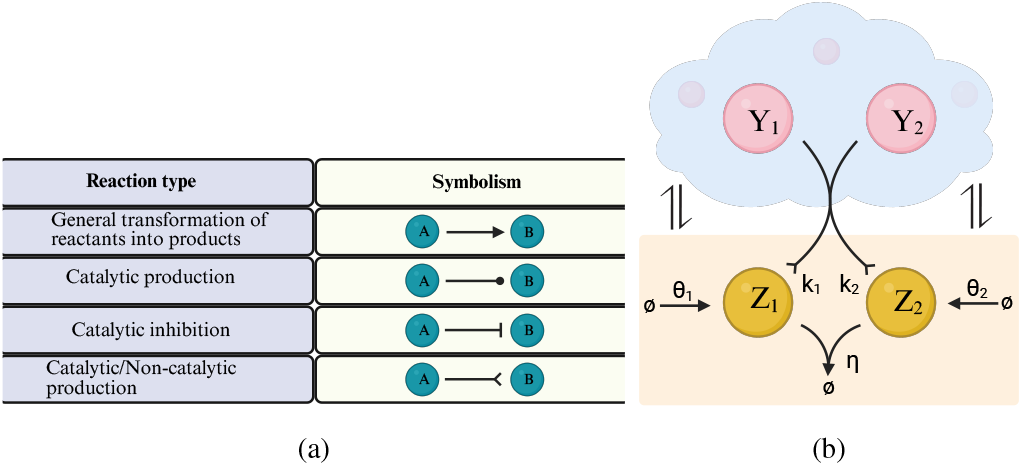
Control design principles. (a) Graphical representation of elementary reaction types [11]. The last symbol denotes either catalytic or non-catalytic production (not a new reaction type). (b) Minimal closed-loop architecture (see the CRN in Eq. (1)). The plant (of unspecified structure) is depicted within a blue cloud, while the controller is enclosed in a beige box. Harpoon arrows denote a communication channel involving an arbitrary number of reactions of arbitrary type through which actuation is achieved.

## III. Minimal Control Design

We present a control architecture composed of two species—the minimum number required—that is capable of driving the product of two species concentrations in a biomolecular process of interest to a prescribed level (setpoint). Low-complexity CRNs are particularly relevant for practical applications, as they offer significant advantages from an experimental standpoint.

We assume a poorly characterized, uncertain process to be controlled (plant) with *q* species, *Y*_1_, *Y*_2_, …, *Y*_*q*−1_, *Y*_*q*_, where *Y*_1_, *Y*_2_ are designated as the target (output) species. We introduce a controller with species *Z*_1_, *Z*_2_, participating in the following chemical reactions (see Fig. 1 b):

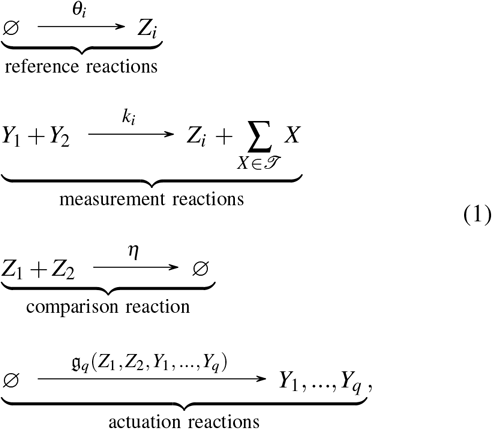

where 𝒯 ⊆ {*Y*_1_,*Y*_2_} and *θ*_*i*_, *k*_*i*_ ∈ ℝ_≥0_ with *i* ∈ {1, 2} and *θ*_1_ + *θ*_2_ > 0, *k*_1_ + *k*_2_ > 0, *θ*_1_ + *k*_1_ > 0, *θ*_2_ + *k*_2_ > 0 and (*θ*_1_ − *θ*_2_)(*k*_2_ − *k*_1_) > 0. These inequalities ensure the presence of at least one reference and one measurement reaction, that each controller species participates in at least one of these reactions, and that the steady state of interest is biologically meaningful. Applying the law of mass action and noting that the reference reaction does not directly affect *Z*_1_, *Z*_2_, we obtain the controller dynamics: 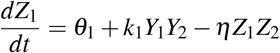 and 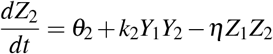. Taking the difference yields the evolution of a non-physical memory variable, enabling integral control action. Specifically, we have

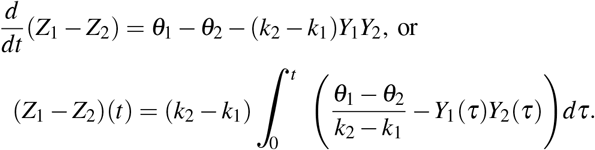

The integrand can be seen as a control error signal where *Y*_1_*Y*_2_ is the output of interest and 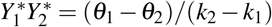 the setpoint (the superscript (·)^∗^ denotes steady-state values). Setpoint tracking is achieved when the error signal converges to zero, which necessitates closed-loop asymptotic stability. However, this property is not always guaranteed for arbitrary network configurations.

As can be seen from the CRN in Eq. (1), we allow the measurement reaction to be disruptive. In particular, when this reaction corresponds to a fully catalytic production process (i.e., *X*_1_ and *X*_2_ are not consumed), output sensing can be achieved without introducing disturbances to the plant, which is the ideal scenario. However, to enhance design flexibility, we also consider scenarios in which this condition cannot be satisfied and loading effects arise due to the measurement process; that is, *X*_1_ or *X*_2_ are consumed. Such effects arising from non-ideal sensing, among others, are common in practical control engineering across many fields, since the implementation of feedback control loops typically requires the interconnection of subsystems that do not exhibit perfect modularity [22], [23]. Moreover, it is evident that the actuation reactions (see the CRN in Eq. (1)) apply the corrective control action on the plant and do not directly affect the controller dynamics. In principle, this actuation mechanism can be implemented via an arbitrary number of reactions of any type (not necessarily obeying mass-action kinetics), provided that the resulting closed-loop system admits a (biologically) meaningful and asymptotically stable steady state. It is worth noting that the control architecture discussed in Section 2.1 of [11] constitutes a special case of the CRN in Eq. (1), involving one reference and one measurement reaction, together with one actuation reaction. The latter is implemented via catalytic inhibition of one of the target species by a controller species.

For a given open-loop network, achieving closed-loop asymptotic stability is a decisive factor for the implementability of the control topologies belonging to the controller family defined by the CRN in Eq. (1). For instance, the following ODE model describing a closed-loop system with two mutually activating target species is unstable for any realistic parameter set (see also Section V):

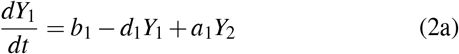

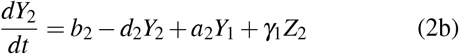

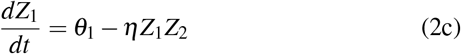

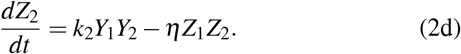

## IV. Closed-Loop Stability Analysis

We first focus on a family of closed-loop systems satisfying the following assumptions:

**S1**. The plant consists of two (target) species, *Y*_1_ and *Y*_2_, and an arbitrary number of reactions of arbitrary type.

**S2**. There is a single reference, measurement and actuation reaction, as defined in the CRN given by Eq. (1). The latter is realized via one target species and one controller species.

This family of closed-loop systems can thus be described as follows

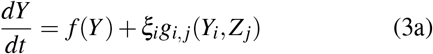

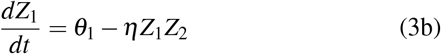

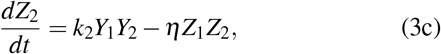

where *i, j* ∈ {1, 2} indexes the family of closed-loop systems, *f* : ℝ^2^ → ℝ^2^, *g*_*i, j*_ : ℝ × ℝ → ℝ, 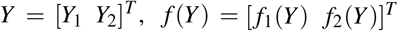, *ξ*_1_ = [1 0]^*T*^, *ξ*_2_ = [0 1]^*T*^. Without loss of generality, we assume that the reference and measurement reactions are carried out via species *Z*_1_ and *Z*_2_, respectively. Note that *f* and *g*_*i, j*_, which may not follow mass-action kinetics, distinguish intrinsic plant dynamics from actuation, enabling a modular interconnection view consistent with Section III.

The Jacobian matrix, evaluated at a steady state of interest 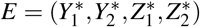, is

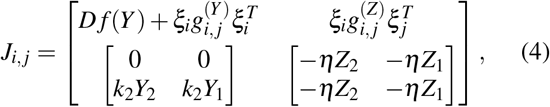

where 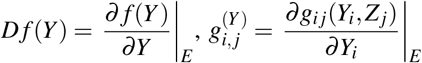 and 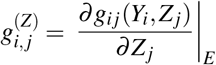.

Eq. (4) can be written as

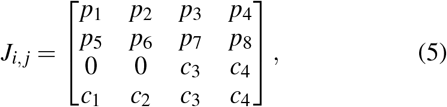

where

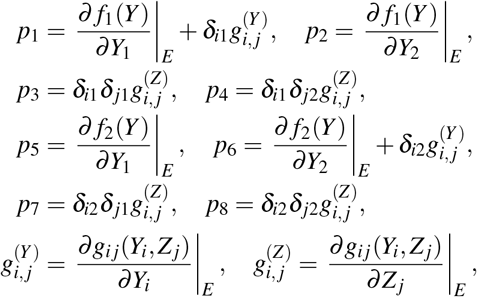

with 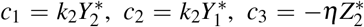, and 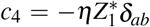 denoting the Kronecker delta, *p*_1_, …, *p*_8_ ∈ ℝ, *c*_1_, *c*_2_ > 0 and *c*_3_, *c*_4_ < 0. Note that *p*_1_, …, *p*_8_ and *c*_1_, *c*_2_ are always independent of *η*.

### Theorem 1.

Consider the steady state 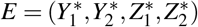 of the system described by Eqs. (3) with the associated Jacobian matrix given by Eq. (5). If one of the following holds: (i) *p*_3_(*c*_1_ *p*_6_ − *c*_2_ *p*_5_) > 0, (ii) *p*_4_(*c*_2_ *p*_5_ − *c*_1_ *p*_6_) > 0, (iii) *p*_7_(*c*_2_ *p*_1_ − *c*_1_ *p*_2_) > 0, or (iv) *p*_8_(*c*_1_ *p*_2_ − *c*_2_ *p*_1_) > 0, then the steady state *E* is unstable.

*Proof:* If det *J*_*i, j*_ < 0, then the 4 × 4 matrix *J*_*i, j*_ has at least one eigenvalue with positive real part; hence, *E* is an unstable steady state of the system described by Eqs. (3).

Note also that:

- *J*_*i, j*_ has four eigenvalues (counting multiplicities).
- The product of the eigenvalues of *J*_*i, j*_ equals det *J*_*i, j*_.
- All entries of *J*_*i, j*_ are real; therefore, any complex eigenvalues occur in conjugate pairs.
- The product of a complex number and its conjugate is real and non-negative.
- Setting the derivatives in Eqs. (3b)–(3c) to zero yields 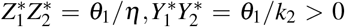, so the steady state 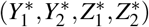 is positive.

We now examine all possible forms of det *J*_*i, j*_:

- For *i* = *j* = 1, we get

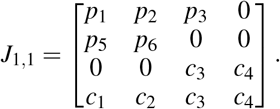 We compute: det*J*_1,1_ = −*c*_2_*c*_4_ *p*_3_ *p*_5_ + *c*_1_*c*_4_ *p*_3_ *p*_6_. If *p*_3_(*c*_1_ *p*_6_ − *c*_2_ *p*_5_) > 0 (condition (i) is true), then det*J*_1,1_ < 0.
- For *i* = 1, *j* = 2, we get

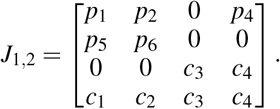 We compute: det*J*_1,2_ = *c*_2_*c*_3_ *p*_4_ *p*_5_ − *c*_1_*c*_3_ *p*_4_ *p*_6_. If *p*_4_(*c*_2_ *p*_5_ − *c*_1_ *p*_6_) > 0 (condition (ii) is true), then det*J*_1,2_ < 0.
- For *i* = 2, *j* = 1, we get

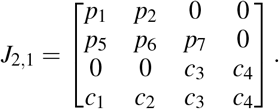 We compute: det*J*_2,1_ = *c*_2_*c*_4_ *p*_1_ *p*_7_ − *c*_1_*c*_4_ *p*_2_ *p*_7_. If *p*_7_(*c*_2_ *p*_1_ − *c*_1_ *p*_2_) > 0 (condition (iii) is true), then det*J*_2,1_ < 0.
- For *i* = *j* = 2, we get

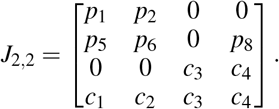 We compute: det*J*_2,2_ = −*c*_2_*c*_3_ *p*_1_ *p*_8_ + *c*_1_*c*_3_ *p*_2_ *p*_8_. If *p*_8_(*c*_1_ *p*_2_ − *c*_2_ *p*_1_) > 0 (condition (iv) is true), then det*J*_2,2_ < 0.

We now introduce the following condition:

**C1**. Assume (*p*_1_ + *p*_6_) < 0 and that one of the following is true:

i. (*p*_1_ + *p*_6_)(*p*_2_ *p*_5_ − *p*_1_ *p*_6_ − *c*_1_ *p*_3_) > *p*_3_(*c*_2_ *p*_5_ − *c*_1_ *p*_6_) > 0,
ii. (*p*_1_ + *p*_6_)(*p*_2_ *p*_5_ − *p*_1_ *p*_6_ + *c*_1_ *p*_4_) > *p*_4_(*c*_1_ *p*_6_ − *c*_2_ *p*_5_) > 0,
iii. (*p*_1_ + *p*_6_)(*p*_2_ *p*_5_ − *p*_1_ *p*_6_ − *c*_2_ *p*_7_) > *p*_7_(*c*_1_ *p*_2_ − *c*_2_ *p*_1_) > 0,
iv. (*p*_1_ + *p*_6_)(*p*_2_ *p*_5_ − *p*_1_ *p*_6_ + *c*_2_ *p*_8_) > *p*_8_(*c*_2_ *p*_1_ − *c*_1_ *p*_2_) > 0.

### Theorem 2.

Consider the steady state 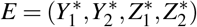 of the system described by Eqs. (3) with the associated Jacobian matrix given by Eq. (5). If condition **C1** is satisfied, then there exists a sufficiently large *η* such that the steady state *E* is locally exponentially stable.

*Proof:* We examine the system described by Eqs. (3) and its Jacobian matrix given by Eq. (5) for all the different pairs of *i, j* ∈ {1, 2}:

- For *i* = *j* = 1, we get

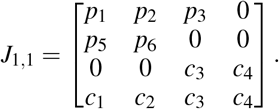 The characteristic polynomial is:

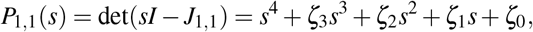

where: *ζ*_3_ = −*c*_3_ − *c*_4_ − *p*_1_ − *p*_6_, *ζ*_2_ = −*p*_2_ *p*_5_ + *p*_1_ *p*_6_ + *c*_3_(*p*_1_ + *p*_6_) + *c*_4_(*p*_1_ + *p*_6_), *ζ*_1_ = *c*_3_(*p*_2_ *p*_5_ − *p*_1_ *p*_6_) + *c*_4_(*p*_2_ *p*_5_ − *p*_1_ *p*_6_ − *c*_1_ *p*_3_), *ζ*_0_ = *c*_4_ *p*_3_(−*c*_2_ *p*_5_ + *c*_1_ *p*_6_) and *I* is the identity matrix of appropriate dimension. By setting the derivatives in Eqs. (3) to zero, we obtain 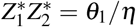 and 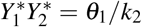, and we also observe that 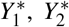 are independent of *η*. These results hold regardless of the values of *i* and *j*. When *i* = *j* = 1, Eq. (3a) shows that 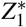 is independent of *η*, too. This is not the case for 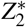, for which we have 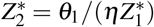. Moreover, *p*_1_, *p*_2_, *p*_3_, *p*_5_, *p*_6_ are independent of *η*. The coefficients of *P*_1,1_(*s*) can be written as

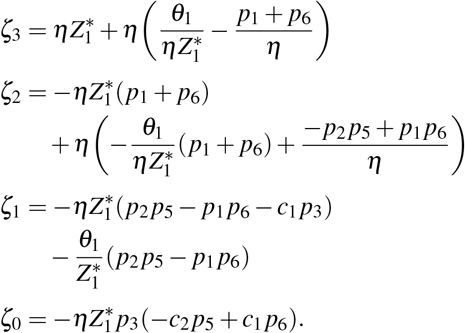 According to the Routh–Hurwitz criterion, *P*_1,1_(*s*) has all its roots in the open left half-plane if and only if *ζ*_3_, *ζ*_2_, *ζ*_1_, *ζ*_0_ > 0, *ζ*_2_*ζ*_3_ > *ζ*_1_ and 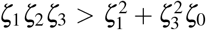. This further guarantees that *E* is a locally exponentially stable steady state of the system described by Eqs. (3). For a sufficiently large *η*, we have *ζ*_3_ > 0, *ζ*_2_ > 0 provided (*p*_1_ + *p*_6_) < 0, *ζ*_1_ > 0 provided (*p*_2_ *p*_5_ − *p*_1_ *p*_6_ − *c*_1_ *p*_3_) < 0, and *ζ*_0_ > 0 provided *p*_3_(−*c*_2_ *p*_5_ + *c*_1_ *p*_6_) < 0. Taking these constraints into account, the inequality *ζ*_2_*ζ*_3_ > *ζ*_1_ also holds. If we additionally assume

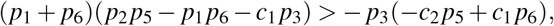

then 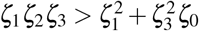 holds as well. Gathering together the above inequalities we arrive at condition **C1**-(i). We follow a similar procedure for the remaining cases.
- For *i* = 1, *j* = 2, we get

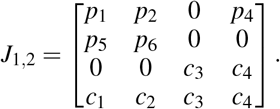 The characteristic polynomial is:

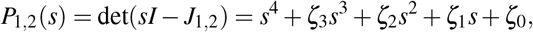

where: *ζ*_3_ = −*c*_3_ − *c*_4_ − *p*_1_ − *p*_6_, *ζ*_2_ = −*c*_1_ *p*_4_ − *p*_2_ *p*_5_ + *p*_1_ *p*_6_ + *c*_3_(*p*_1_ + *p*_6_) + *c*_4_(*p*_1_ + *p*_6_), *ζ*_1_ = *c*_4_(*p*_2_ *p*_5_ − *p*_1_ *p*_6_) + *c*_3_(*p*_2_ *p*_5_ − *p*_1_ *p*_6_ + *c*_1_ *p*_4_) − *c*_2_ *p*_4_ *p*_5_ + *c*_1_ *p*_4_ *p*_6_, *ζ*_0_ = *c*_3_ *p*_4_(*c*_2_ *p*_5_ − *c*_1_ *p*_6_). From Eqs. (3) we can see that 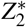 is independent of *η* and 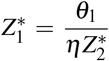. Moreover, *p*_1_, *p*_2_, *p*_4_, *p*_5_, *p*_6_ are independent of *η*. The coefficients of *P*_1,2_(*s*) can be written as

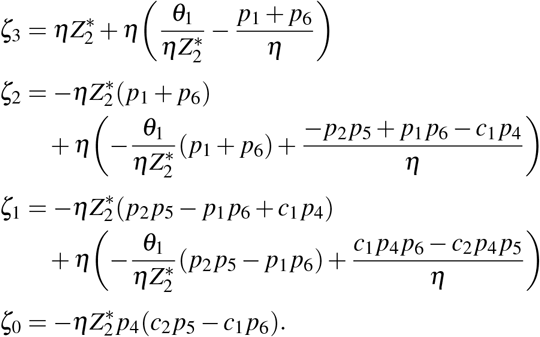 For a sufficiently l arge *η*, we have *ζ* _3_ > 0, *ζ* _2_ > 0 provided (*p*_1_ + *p*_6_) < 0, *ζ*_1_ > 0 provided (*p*_2_ *p*_5_ − *p*_1_ *p*_6_ + *c*_1_ *p*_4_) < 0 and *ζ*_0_ > 0 provided *p*_4_(*c*_2_ *p*_5_ − *c*_1_ *p*_6_) < 0. Taking these constraints into account, the inequality *ζ*_2_*ζ*_3_ > *ζ*_1_ also holds. If we additionally assume

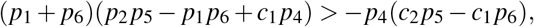

then 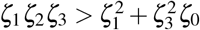 holds as well. Gathering together the above inequalities we arrive at condition **C1**-(ii).
- For *i* = 2, *j* = 1, we get

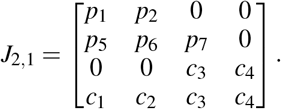 The characteristic polynomial is:

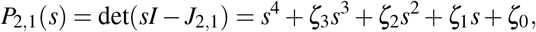

where: *ζ*_3_ = −*c*_3_ − *c*_4_ − *p*_1_ − *p*_6_, *ζ*_2_ = −*p*_2_ *p*_5_ + *p*_1_ *p*_6_ + *c*_3_(*p*_1_ + *p*_6_) + *c*_4_(*p*_1_ + *p*_6_), *ζ*_1_ = *c*_3_(*p*_2_ *p*_5_ − *p*_1_ *p*_6_) + *c*_4_(*p*_2_ *p*_5_ − *p*_1_ *p*_6_ − *c*_2_ *p*_7_), *ζ*_0_ = *c*_4_ *p*_7_(*c*_2_ *p*_1_ − *c*_1_ *p*_2_). From Eqs. (3) we can see that 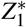 is independent of *η* and 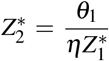. Moreover, *p*_1_, *p*_2_, *p*_5_, *p*_6_, *p*_7_ are independent of *η*. The coefficients of *P*_2,1_(*s*) can be written as

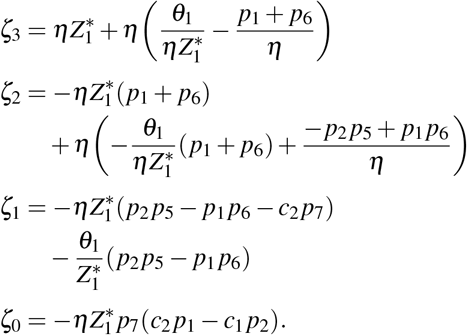 For a sufficiently large *η*, we have *ζ*_3_ > 0, *ζ*_2_ > 0 provided (*p*_1_ + *p*_6_) < 0, *ζ*_1_ > 0 provided (*p*_2_ *p*_5_ − *p*_1_ *p*_6_ − *c*_2_ *p*_7_) < 0 and *ζ*_0_ > 0 provided *p*_7_(*c*_2_ *p*_1_ − *c*_1_ *p*_2_) < 0. Taking these constraints into account, the inequality *ζ*_2_*ζ*_3_ > *ζ*_1_ also holds. If we additionally assume

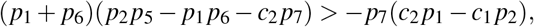

then 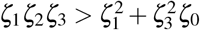 holds as well. Gathering together the above inequalities we arrive at condition **C1**-(iii).
- For *i* = 2, *j* = 2, we get

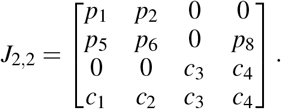 The characteristic polynomial is:

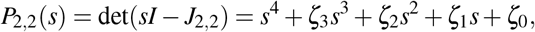

where: *ζ*_3_ = −*c*_3_ − *c*_4_ − *p*_1_ − *p*_6_, *ζ*_2_ = −*c*_2_ *p*_8_ − *p*_2_ *p*_5_ + *p*_1_ *p*_6_ + *c*_3_(*p*_1_ + *p*_6_) + *c*_4_(*p*_1_ + *p*_6_), *ζ*_1_ = *c*_4_(*p*_2_ *p*_5_ − *p*_1_ *p*_6_) + *c*_3_(*p*_2_ *p*_5_ − *p*_1_ *p*_6_ + *c*_2_ *p*_8_) + *c*_2_ *p*_1_ *p*_8_ − *c*_1_ *p*_2_ *p*_8_, *ζ*_0_ = *c*_3_ *p*_8_(*c*_1_ *p*_2_ − *c*_2_ *p*_1_). From Eqs. (3) we can see that 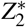 is independent of *η* and 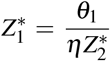. Moreover, *p*_1_, *p*_2_, *p*_5_, *p*_6_, *p*_8_ are independent of *η*. The coefficients of *P*_2,2_(*s*) can be written as

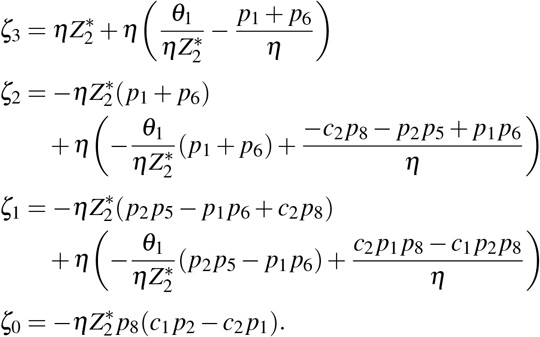 For a sufficiently large *η*, we have *ζ*_3_ > 0, *ζ*_2_ > 0 provided (*p*_1_ + *p*_6_) < 0, *ζ*_1_ > 0 provided (*p*_2_ *p*_5_ − *p*_1_ *p*_6_ + *c*_2_ *p*_8_) < 0 and *ζ*_0_ > 0 provided *p*_8_(*c*_1_ *p*_2_ − *c*_2_ *p*_1_) < 0. Taking these constraints into account, the inequality *ζ*_2_*ζ*_3_ > *ζ*_1_ also holds. If we additionally assume

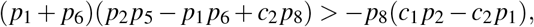

then 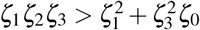 holds as well. Gathering together the above inequalities we arrive at condition **C1**-(iv).

**Theorem 1** provides sufficient conditions for instability that can be used to rule out architectures that are not feasible. Note that, to satisfy these conditions, it is not always necessary to know the parameter regimes of interest—the structural characteristics of the closed-loop system may suffice. On the other hand, **Theorem 2** provides sufficient conditions for local exponential stability of an equilibrium of interest. These conditions are based on the so-called *fast sequestration regime* [24], [25], a common assumption that allows *η* to be made arbitrarily large. Note also that each theorem relies on a (different) set of four conditions, each corresponding to a different actuation mechanism, realized through species (i) *Y*_1_, *Z*_1_, (ii) *Y*_1_, *Z*_2_, (iii) *Y*_2_, *Z*_1_, and (iv) *Y*_2_, *Z*_2_.

We now move to a more general family of closed-loop systems by relaxing the assumptions S1, S2, specifically by replacing them with the following:

**S1**′. The plant includes *q* > 3 species and reactions of arbitrary type, while *Y*_1_ and *Y*_2_ are the target species.

**S2**′. There is one reference, one measurement, and one actuation reaction, as defined by the CRN in Eq. (1). The latter is realized via one target species and one controller species while other (non-target) species from the plant are allowed to participate. Both controller species may also participate as catalysts (without being consumed) in an arbitrary number of reactions of arbitrary type that do not include the target species as reactants or products.

The resulting family of closed-loop systems can be written as follows

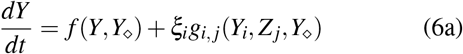

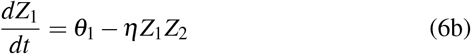

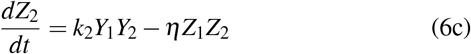

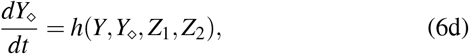

where *i, j* ∈ {1, 2} indexes the family of closed-loop systems, *f* : ℝ^*q*^ → ℝ^2^, *g*_*i, j*_ : ℝ × ℝ → ℝ, *h* : ℝ^*q*+2^ → ℝ^*q*−2^, *Y* = [*Y*_1_ *Y*_2_]^*T*^, *f*(*Y,Y*_⋄_) = [*f*_1_(*Y,Y*_⋄_) *f*_2_(*Y,Y*_⋄_)]^*T*^, *ξ* = [1 0]^*T*^, *ξ*_2_ = [0 1]^*T*^, *Y*_⋄_ = [*Y*_3_ *Y*_4_ … *Y*_*q*_]^*T*^. Moreover, *h* = [*h*_1_ … *h*_*q*−2_]^*T*^, *h*_*ℓ*_ : ℝ^*q*+2^ → ℝ, so that 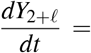 *h*_*ℓ*_(*Y,Y*_⋄_, *Z*_1_, *Z*_2_), *ℓ* = 1, …, *q* − 2. Also, *f, g*_*i, j*_, *h* are permitted to follow non–mass-action kinetics.

The Jacobian matrix, evaluated at a steady state of interest 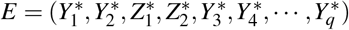 is

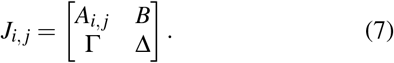

*A*_*i, j*_ has the same structure as the matrix given by Eq. (4) and can be expressed as

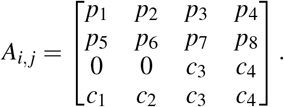

where

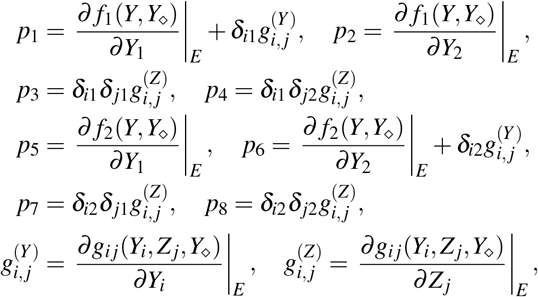

with 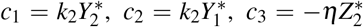, and 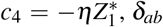 denoting the Kronecker delta, *p*_1_, …, *p*_8_ ∈ ℝ, *c*_1_, *c*_2_ > 0 and *c*_3_, *c*_4_ < 0. The remaining matrices *B* ∈ ℝ^4×(*q*−2)^, Γ ∈ ℝ^(*q*−2)×4^ and Δ ∈ ℝ^(*q*−2)×(*q*−2)^, respectively, are defined entrywise as

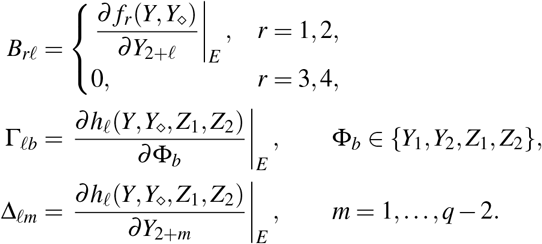

### Theorem 3.

Consider the steady state 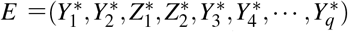 of the system described by Eqs. (6) with associated Jacobian matrix given by Eq. (7). Assume that (a) Δ is Hurwitz, (b) Condition **C1** holds, and (c) one of the following holds: (i) *B* = 0, (ii) Γ = 0, (iii)^1^ 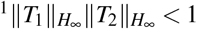, where *T*_1_(*s*) = (*sI* − *A*_*i, j*_)^−1^*B, T*_2_(*s*) = (*sI* − Δ)^−1^Γ, *s* denotes the Laplace variable, and *I* denotes the identity matrix of appropriate dimension. Then there exists a sufficiently large *η* such that the steady state *E* is locally exponentially stable.

*Proof:* If condition (b) is satisfied, then there exists a sufficiently large *η* such that *A*_*i, j*_ is Hurwitz. This follows from the proof of **Theorem 2**, where we establish the same result for a matrix with the same structure as *A*_*i, j*_, namely the matrix given in Eq. (4).

If *B* = 0 (condition (c)-(i) is true), *J*_*i, j*_ given by Eq. (7) becomes

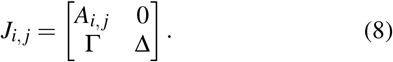

Eq. (8) is a block lower triangular matrix. Since both *A*_*i, j*_ and Δ (see condition (a)) are Hurwitz, then *J*_*i, j*_ is Hurwitz. Similarly, for Γ = 0 (condition (c)-(ii) is true), we have

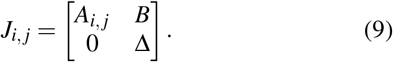

Eq. (9) is a block upper triangular matrix which is Hurwitz because both *A*_*i, j*_ and Δ are Hurwitz.

Finally, around the steady state *E*, the system described by Eqs. (6) can be seen as the pure feedback interconnection of two subsystems, i.e.:

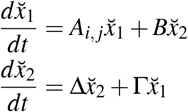

where 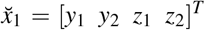 and 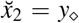 with 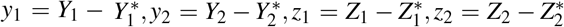 and 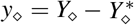 representing sufficiently small perturbations around the steady state *E*.

Both subsystems are asymptotically stable as *A*_*i, j*_, Δ are Hurwitz. We can further compute the corresponding transfer functions as *T*_1_(*s*) = (*sI* − *A*_*i, j*_)^−1^*B* and *T*_2_(*s*) = (*sI* − Δ)^−1^Γ.

By the Small-Gain Theorem (see Corollary 3.1 in [26]), if

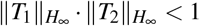

(condition (c)-(iii) is true), then the pure feedback interconnection of the two subsystems is asymptotically stable.

Consequently, if conditions (a)–(c) hold, there exists a sufficiently large *η* such that *E* is a locally exponentially stable steady state for the system described by Eqs. (6).

**Theorem 3** extends the results of **Theorem 2** by providing sufficient conditions for the local exponential stability of an equilibrium of interest within a more general family of closed-loop systems. Note that *B* = 0 implies that species *Y*_3_,*Y*_4_, *· · ·, Y*_*q*_ do not affect the production or inhibition of species *Y*_1_,*Y*_2_, *Z*_1_, *Z*_2_, while the reverse is implied in case Γ = 0.

## V. Demonstrative Application

Fig. 2 a depicts a closed-loop system with a plant composed of two target species, *Y*_1_ and *Y*_2_, each undergoing a generic birth–death process while also catalyzing the production of the other. This plant was previously studied in [11], where more details about its dynamics in isolation can be found. The controller follows the architecture described by the CRN in Eq. (1) and is capable of robustly driving the product of the target species concentrations, *Y*_1_*Y*_2_, to a setpoint (see Fig. 2 b). In particular, the closed-loop dynamics can be described by

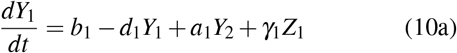

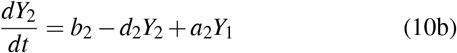

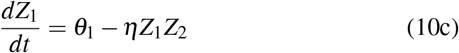

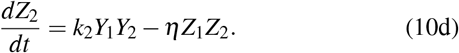

**Fig. 2:**
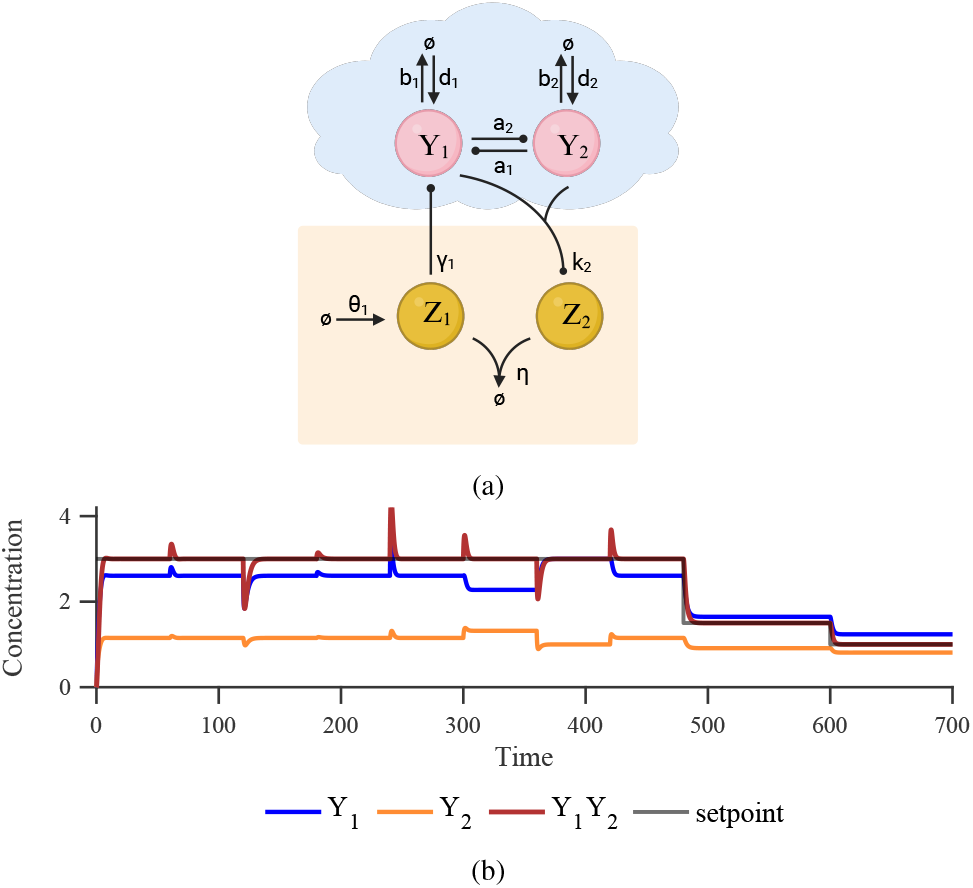
Regulation of the product of two species concentrations. (a) Closed-loop architecture where *Y*_1_ and *Y*_2_ are the target species and *Y*_1_*Y*_2_ is the output quantity of interest. (b) Simulated response of the ODE model in Eq. (10) using the nominal parameter values *b*_1_ = *b*_2_ = 1, *d*_1_ = *d*_2_ = 2, *a*_1_ = *a*_2_ = 0.5, *θ*_1_ = 3, *k*_2_ = 1, *γ*_1_ = 0.5, and *η* = 1000. Every 60 time units, a *±*50% parameter perturbation is applied in the following order (arbitrarily chosen): *b*_1_, *d*_1_, *a*_1_, *γ*_1_, *b*_2_, *d*_2_, *a*_2_, *θ*_1_, *η*, and *k*_2_. Notably, only changes in *θ*_1_, *k*_2_ affect 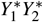, despite other parameter changes altering 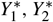 individually.

Following the formalism adopted in Section IV, the Jacobian matrix, evaluated at a steady state of interest 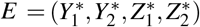, can be expressed in the form of Eq. (5):

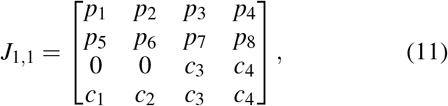

where 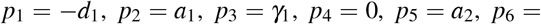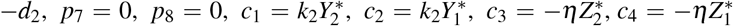. As can be seen by setting the derivatives in Eqs. (10c), (10d) to zero, 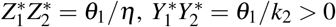. The system given by Eqs. (10) satisfies assumptions **S1, S2**. Consequently, by applying **Theorem 2** we conclude that if 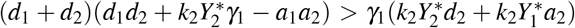, then there exists a sufficiently large *η* such that *E* is locally exponentially stable. Note that **Theorem 1** can also be informative from a control-design perspective here. For example, for the plant under consideration, it would be im-possible—regardless of the parameter regime—to implement the same actuation mechanism between species *Y*_2_ and *Z*_2_, i.e., 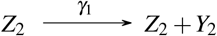 (see also Eqs (2)). The reason is that the entries of the corresponding Jacobian, *J*_2,2_, would be identical to those of *J*_1,1_, except that *p*_3_ = 0 and *p*_8_ = *γ*_1_ and, thus, *p*_8_(*c*_1_ *p*_2_ − *c*_2_ *p*_1_) > 0 would always be true. According to **Theorem 1**, this condition entails instability.

We now consider a third birth–death process involving species *Y*_3_, which interacts with the above system via catalytic production reactions, as illustrated in Fig. 3 a. The dynamics of the overall system can be described by

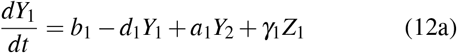

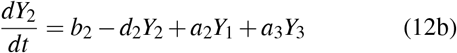

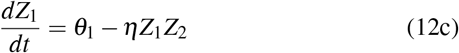

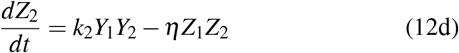

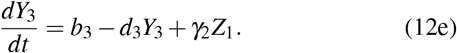

**Fig. 3:**
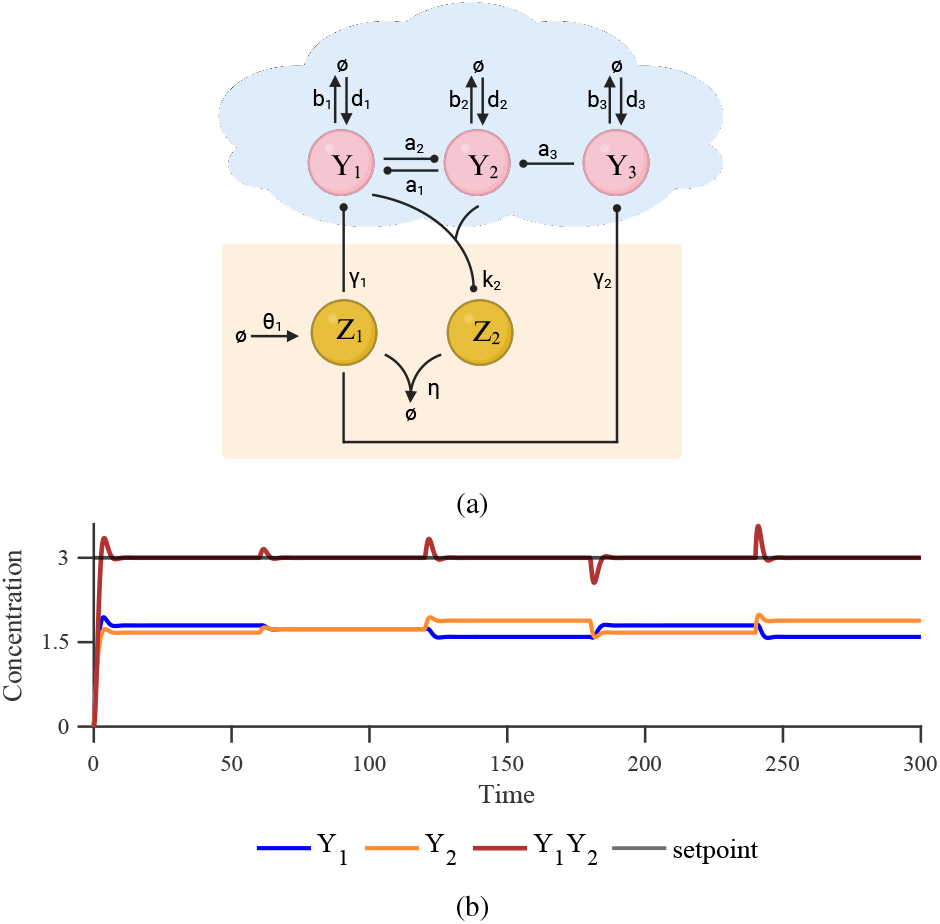
Regulation of the product of two species concentrations in the presence of a third species. (a) Closed-loop architecture obtained by interconnecting the architecture in Fig. 2 a with an additional one-species process. The target species and the output quantity of interest remain unchanged. (b) Simulated response of the ODE model in Eq. (12) using the parameter values *b*_3_ = 4, *d*_3_ = 2, *a*_3_ = 0.5, and *γ*_2_ = 0.5, with all remaining parameter values identical to those in Fig. 2 b. Every 60 time units, a ±50% parameter perturbation is applied in the following order (arbitrarily chosen): *γ*_2_, *b*_3_, *d*_3_, and *a*_3_.

Note that this new system, given by Eqs (12), satisfies assumptions **S1**′, **S2**′. Also, *Y*_1_*Y*_2_ remains the output quantity of interest. The corresponding Jacobian matrix, evaluated at a steady state of interest 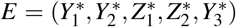, can be expressed in the form of Eq. (7) as

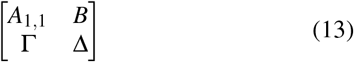

where *A*_1,1_ is given by Eq. (11), *B* = [0 *a*_3_ 0 0]^*T*^, Γ = [0 0 *γ*_2_ 0], Δ = [−*d*_3_]. We now invoke **Theorem 3** to study the overall stability. Condition (a) is immediately satisfied. Moreover, if we additionally assume that 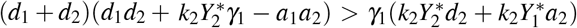, then condition (b) holds as well. Finally, condition (c) can be ensured by considering the special case *a*_3_ = 0 or *γ*_2_ = 0, while, in the more general case where *a*_3_, *γ*_2_ > 0, by requiring 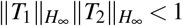 Therefore, we can find a sufficiently large *η* such that *E* is locally exponentially stable. In Fig. 3 b, we demonstrate the robust steady-state tracking capability of the system described by Eqs. (12). Note that, given the nominal parameter values used in Fig. 2, overall stability can be guaranteed under the constraint *γ*_2_*a*_3_ < 0.33*d*_3_.

## VI. A Generalized Control Formulation

The control architecture of Section III can be extended to accommodate an arbitrary number of target and controller species and a broader class of control objectives.

Consider a poorly characterized and uncertain plant consisting of *q* species *Y*_1_,*Y*_2_, …, *Y*_*q*_, of which *n* species *Y*_1_,*Y*_2_, …, *Y*_*n*_ are target species (*q* ≥ *n*), and a controller with *φ* species *Z*_1_, *Z*_2_, …, *Z*_*φ*_. Our goal is to drive a prescribed monomial of the target species concentrations, namely 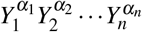 with *α*_1_, …, *α*_*n*_ ∈ ℤ_>0_, to a setpoint. Before introducing the general mathematical formulation, we first illustrate the proposed idea through a simple example. In Fig. 4, we build upon the case study of Fig. 3 and modify the control architecture by introducing two additional species. This allows us to achieve robust steady-state tracking with respect to 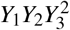 (output quantity of interest). The dynamics of the resulting closed-loop system can be described by

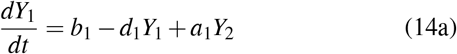

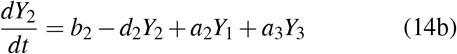

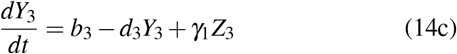

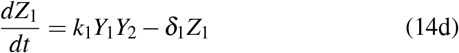

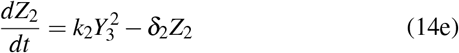

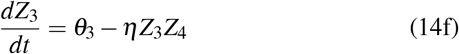

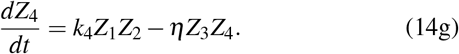

**Fig. 4:**
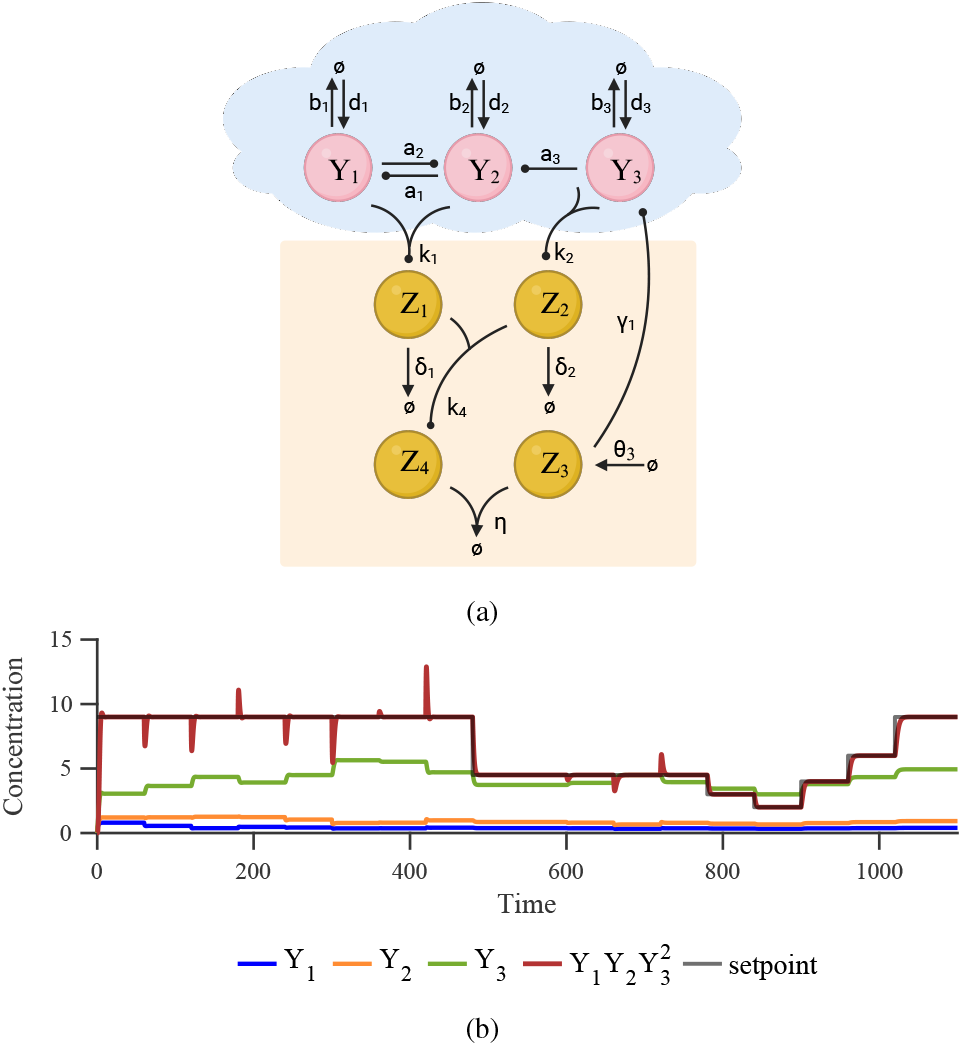
Regulation of a monomial of three species concentrations. (a) Closed-loop architecture where *Y*_1_, *Y*_2_, and *Y*_3_ are the target species and 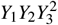 is the output quantity of interest. The plant has the same structure as in Fig. 3 a. (b) Simulated response of the ODE model in Eq. (14) using the nominal parameter values *b*_1_ = *b*_2_ = *b*_3_ = 1, *d*_1_ = *d*_2_ = *d*_3_ = 2, *a*_1_ = *a*_2_ = *a*_3_ = 0.5, *γ*_1_ = 1, *θ*_3_ = 1, *k*_1_ = *k*_2_ = *k*_4_ = 1, *δ*_1_ = *δ*_2_ = 3, and *η* = 1000. Every 60 time units, a *±*50% parameter perturbation is applied in the following order (arbitrarily chosen): *b*_1_, *d*_1_, *a*_1_, *b*_2_, *d*_2_, *a*_2_, *a*_3_, *θ*_3_, *η, b*_3_, *d*_3_, *γ*_1_, *k*_2_, *k*_1_, *k*_4_, *δ*_1_, and *δ*_2_.

Eqs. (14d)–(14g) yield, at steady state,

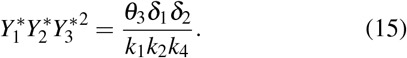

The above control architecture can be viewed as a special case of a more general control framework consisting of the following groups of chemical reactions (see Fig. 5):

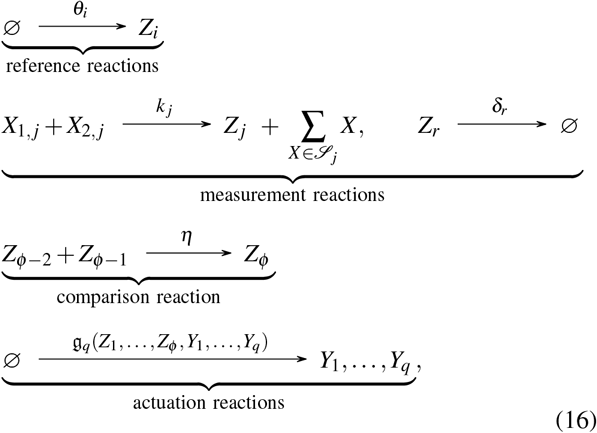

where *i* ∈ {*φ* − 2, *φ* − 1}, *j* ∈ {1,…, *φ* − 1}, and *r* ∈ {1,…, *φ*} \ {*φ* − 2, *φ* − 1}, with *θ*_*i*_, *k* _*j*_ ∈ ℝ_≥0_ and 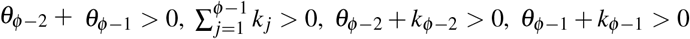, assuming finite and positive steady states of interest. These constraints are analogous to those associated with Eq. (1). The degradation rate *δ*_*φ*_ may depend on other species; that is, *δ*_*φ*_ = *δ* (*Y*_1_, …, *Y*_*q*_, *Z*_1_, …, *Z*_*φ*−1_) (whereas *δ*_*r*_ is constant for *r* ∈ {1,…, *φ* − 3}). The species involved in the measurement reactions belong to the target set 𝒴 = {*Y*_1_, …, *Y*_*n*_} and the (partial controller) set 𝒵 = {*Z*_1_, …, *Z*_*φ*−3_}. For each production measurement reaction, the reactants satisfy *X*_1, *j*_, *X*_2, *j*_ ∈ 𝒴 ∪ 𝒵, and 𝒮_*j*_ ⊆ 𝒴 ∪ 𝒵 denotes a (possibly empty) set of species produced by the *j*-th reaction. The dependencies among species induced by the measurement reactions are referred to as the *measurement cascade*. We assume that controller species are not consumed by the cascade, i.e., {*X*_1, *j*_, *X*_2, *j*_} ∩ 𝒵 ⊆ 𝒮_*j*_ for *j* = 1, …, *φ* − 1, whereas target species may be consumed. We further assume that every species in 𝒴 ∪ 𝒵 can influence at least one of the terminal species *Z*_*φ*−2_ or *Z*_*φ*−1_ through a finite chain of measurement reactions. Finally, the two terminal measurement reactions producing *Z*_*φ*−2_ and *Z*_*φ*−1_ encode the same monomial of the target species, possibly scaled by different gains (one of which may be zero); that is, the corresponding *terminal measurement monomials* are proportional to 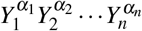.

**Fig. 5:**
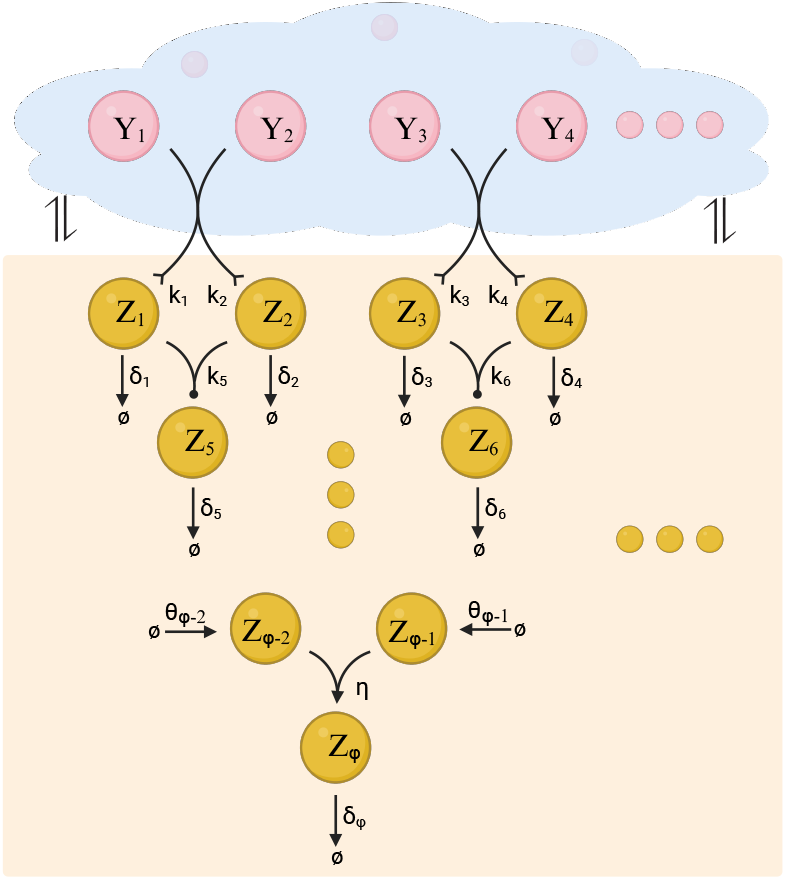
Generalized closed-loop architecture (see the CRN in Eq. (16)). The same visual convention as in Fig. 1b is used. For clarity, only a subset of representative reactions is shown explicitly, while smaller spheres indicate the presence of an arbitrary number of additional reactions and species.

The controller dynamics are therefore

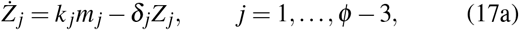

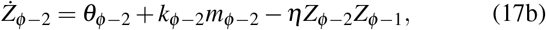

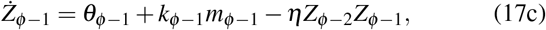

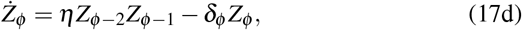

where *m*_*j*_ := *X*_1, *j*_*X*_2, *j*_.

From Eqs. (17b)–(17c) we obtain

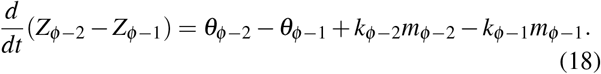

Eqs. (17a) and (18) give, at steady state,

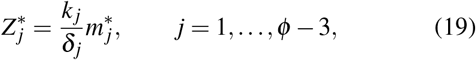

and

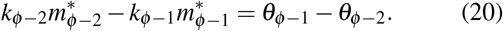

Using Eq. (19) repeatedly to eliminate the controller species in 𝒵 (as dictated by the measurement cascade), the terminal measurement monomials can be expressed in terms of the target steady states. In particular, we assume gains *κ*_*t*2_, *κ*_*t*1_ that depend only on 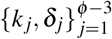 and can be written as products of ratios *k*_*ℓ*_*/δ*_*ℓ*_ ≥ 0. More precisely, there exist integers *ν*_*ℓ,φ*−2_, *ν*_*ℓ,φ*−1_ ∈ ℤ_≥0_ such that

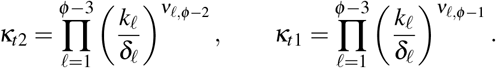

Consequently, the terminal measurement monomials can be written as

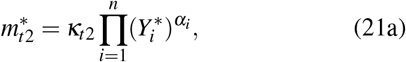

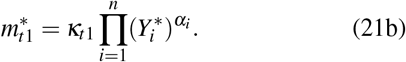

Substituting Eqs. (21) into Eq. (20) gives

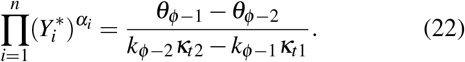

The left-hand side of Eq. (22) represents the output quantity of interest, which is steered to the setpoint given by the right-hand side (we assume (*θ*_*φ*−1_ − *θ*_*φ*−2_)(*k*_*φ*−2_*κ*_*t*2_ − *k*_*φ*−1_*κ*_*t*1_) > 0). The actuation reactions may be of arbitrary type and kinetics, provided the closed-loop system admits a biologically meaningful and asymptotically stable steady state.

## VII. Conclusion

In this work, we investigate the implementation of regulatory topologies for multiplicative control objectives through biomolecular interactions. We first introduce a structurally minimal control architecture comprising two con-troller species. This architecture achieves robust steady-state tracking of an output defined as the product of two target species concentrations. Under biologically meaningful assumptions, we analyze the behavior of two broad families of closed-loop systems of increasing structural complexity and generality. In particular, we derive sufficient conditions for asymptotic stability and instability, which constitute critical design considerations. To corroborate our theoretical analysis, we examine two practical examples: a two-species plant with a two-species controller and a three-species plant with a two-species controller. We then extend the second example with a four-species controller capable of regulating the product of two species concentrations multiplied by the square of a third species concentration. The latter stems from a more general family of control architectures, for which a detailed discussion is provided.

We believe that the theoretical design principles developed herein can motivate the experimental implementation of sophisticated biomolecular control systems. For example, our control architectures are well suited for molecular programming applications. Notably, [11], [20] provide in-depth discussions of DNA strand-displacement realizations of mass-action CRNs with arbitrary structure. It is also experimentally established that within this setting reaction rates can be adjusted over multiple orders of magnitude, enabling operation in the fast sequestration regime (i.e., sufficiently large *η*) [27].

Natural extensions of this work include analyzing basins of attraction and global convergence, identifying critical thresholds for *η* as well as studying stability under slower sequestration regimes, and deriving conditions that preserve setpoint tracking under additional controller species degradation. Another interesting direction is the stochastic analysis of the proposed systems. While deterministic dynamics approximate average behavior at sufficiently high molecule copy numbers, this assumption may not always hold in practice. Such analysis can be carried out computationally, e.g., via *Gillespie’s stochastic simulation algorithm* [28], or analytically using approximation techniques such as moment-closure methods [29].

## Footnotes

1 The *H*_∞_ norm of a matrix *T* (*jω*) is defined as 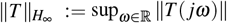 [26].

## References

[1] L. Grozinger, M. Amos, T. E. Gorochowski, P. Carbonell, D. A. Oyarzún, R. Stoof, H. Fellermann, P. Zuliani, H. Tas, and A. Goñi-Moreno, “Pathways to cellular supremacy in biocomputing,” Nature communications, vol. 10, no. 1, p. 5250, 2019.

[2] M. P. McNerney, K. E. Doiron, T. L. Ng, T. Z. Chang, and P. A. Silver, “Theranostic cells: emerging clinical applications of synthetic biology,” Nature Reviews Genetics, vol. 22, no. 11, pp. 730–746, 2021.

[3] N. Sarraf, K. R. Rodriguez, and L. Qian, “Modular reconfiguration of dna origami assemblies using tile displacement,” Science Robotics, vol. 8, no. 77, p. eadf1511, 2023.

[4] C. Gilbert and T. Ellis, “Biological engineered living materials: growing functional materials with genetically programmable properties,” ACS synthetic biology, vol. 8, no. 1, pp. 1–15, 2018.

[5] S. M. Brooks and H. S. Alper, “Applications, challenges, and needs for employing synthetic biology beyond the lab,” Nature Communications, vol. 12, no. 1, p. 1390, 2021.

[6] D. Del Vecchio, Y. Qian, R. M. Murray, and E. D. Sontag, “Future systems and control research in synthetic biology,” Annual Reviews in Control, vol. 45, pp. 5–17, 2018.

[7] M. Khammash, M. Di Bernardo, and D. Di Bernardo, “Cybergenetics: Theory and methods for genetic control systems,” in 2019 IEEE 58th Conference on Decision and Control (CDC). IEEE, 2019, pp. 916–926.

[8] M. Filo, C.-H. Chang, and M. Khammash, “Biomolecular feedback controllers: from theory to applications,” Current Opinion in Biotechnology, vol. 79, p. 102882, 2023.

[9] N. Shakiba, R. D. Jones, R. Weiss, and D. Del Vecchio, “Context-aware synthetic biology by controller design: Engineering the mammalian cell,” Cell systems, vol. 12, no. 6, pp. 561–592, 2021.

[10] E. Alexis, S. Espinel-Ríos, I. G. Kevrekidis, and J. L. Avalos, “Biochemical implementation of acceleration sensing and pida control,” npj Systems Biology and Applications, vol. 11, no. 1, p. 39, 2025.

[11] E. Alexis, C. C. Schulte, L. Cardelli, and A. Papachristodoulou, “Regulation strategies for two-output biomolecular networks,” Journal of the Royal Society Interface, vol. 20, no. 205, p. 20230174, 2023.

[12] A. M. Zand, S. Anastassov, T. Frei, and M. Khammash, “Multi-layer autocatalytic feedback enables integral control amidst resource competition and across scales,” ACS Synthetic Biology, vol. 14, no. 4, pp. 1041–1061, 2025.

[13] K. Stephens, M. Pozo, C.-Y. Tsao, P. Hauk, and W. E. Bentley, “Bacterial co-culture with cell signaling translator and growth controller modules for autonomously regulated culture composition,” Nature communications, vol. 10, no. 1, p. 4129, 2019.

[14] W. Jiang, X. Yang, F. Gu, X. Li, S. Wang, Y. Luo, Q. Qi, and Q. Liang, “Construction of synthetic microbial ecosystems and the regulation of population proportion,” ACS Synthetic Biology, vol. 11, no. 2, pp. 538–546, 2022.

[15] R. D. McCardell, A. Pandey, and R. M. Murray, “Control of density and composition in an engineered two-member bacterial community,” BioRxiv, p. 632174, 2019.

[16] A. Boo, H. Mehta, R. L. Amaro, and G.-B. Stan, “Host-aware rna-based control of synthetic microbial consortia,” Biorxiv, pp. 2023–05, 2023.

[17] U. Kwon, H.-H. Huang, J. L. Chávez, K. Beabout, S. Harbaugh, and D. Del Vecchio, “Incoherent merger network for robust ratiometric gene expression response,” Nucleic Acids Research, vol. 51, no. 6, pp. 2963–2973, 2023.

[18] C. Briat, A. Gupta, and M. Khammash, “Antithetic integral feedback ensures robust perfect adaptation in noisy biomolecular networks,” Cell systems, vol. 2, no. 1, pp. 15–26, 2016.

[19] S. K. Aoki, G. Lillacci, A. Gupta, A. Baumschlager, D. Schweingruber, and M. Khammash, “A universal biomolecular integral feedback controller for robust perfect adaptation,” Nature, vol. 570, no. 7762, pp. 533–537, 2019.

[20] M. Whitby, L. Cardelli, M. Kwiatkowska, L. Laurenti, M. Tribastone, and M. Tschaikowski, “PID control of biochemical reaction networks,” IEEE Transactions on Automatic Control, 2021.

[21] V. Chellaboina, S. P. Bhat, W. M. Haddad, and D. S. Bernstein, “Modeling and analysis of mass-action kinetics,” IEEE Control Systems Magazine, vol. 29, no. 4, pp. 60–78, 2009.

[22] D. Del Vecchio, A. J. Ninfa, and E. D. Sontag, “Modular cell biology: retroactivity and insulation,” Molecular systems biology, vol. 4, no. 1, p. 161, 2008.

[23] J. C. Willems, “The behavioral approach to open and interconnected systems,” IEEE control systems magazine, vol. 27, no. 6, pp. 46–99, 2007.

[24] M. Filo, S. Kumar, and M. Khammash, “A hierarchy of biomolecular proportional-integral-derivative feedback controllers for robust perfect adaptation and dynamic performance,” Nature Communications, vol. 13, no. 1, pp. 1–19, 2022.

[25] N. Olsman, A.-A. Baetica, F. Xiao, Y. P. Leong, R. M. Murray, and J. C. Doyle, “Hard limits and performance tradeoffs in a class of antithetic integral feedback networks,” Cell systems, vol. 9, no. 1, pp. 49–63, 2019.

[26] A. Isidori, Lectures in feedback design for multivariable systems. Springer, 2017.

[27] B. Yurke, A. J. Turberfield, A. P. Mills Jr, F. C. Simmel, and J. L. Neumann, “A dna-fuelled molecular machine made of dna,” Nature, vol. 406, no. 6796, pp. 605–608, 2000.

[28] D. T. Gillespie, “A general method for numerically simulating the stochastic time evolution of coupled chemical reactions,” Journal of computational physics, vol. 22, no. 4, pp. 403–434, 1976.

[29] M. Soltani, C. A. Vargas-Garcia, and A. Singh, “Conditional moment closure schemes for studying stochastic dynamics of genetic circuits,” IEEE transactions on biomedical circuits and systems, vol. 9, no. 4, pp. 518–526, 2015.

